## Supplementary Data for "Kinases in motion: impact of protein and small molecule interactions on kinase conformations"

#### Figure Supplement 1

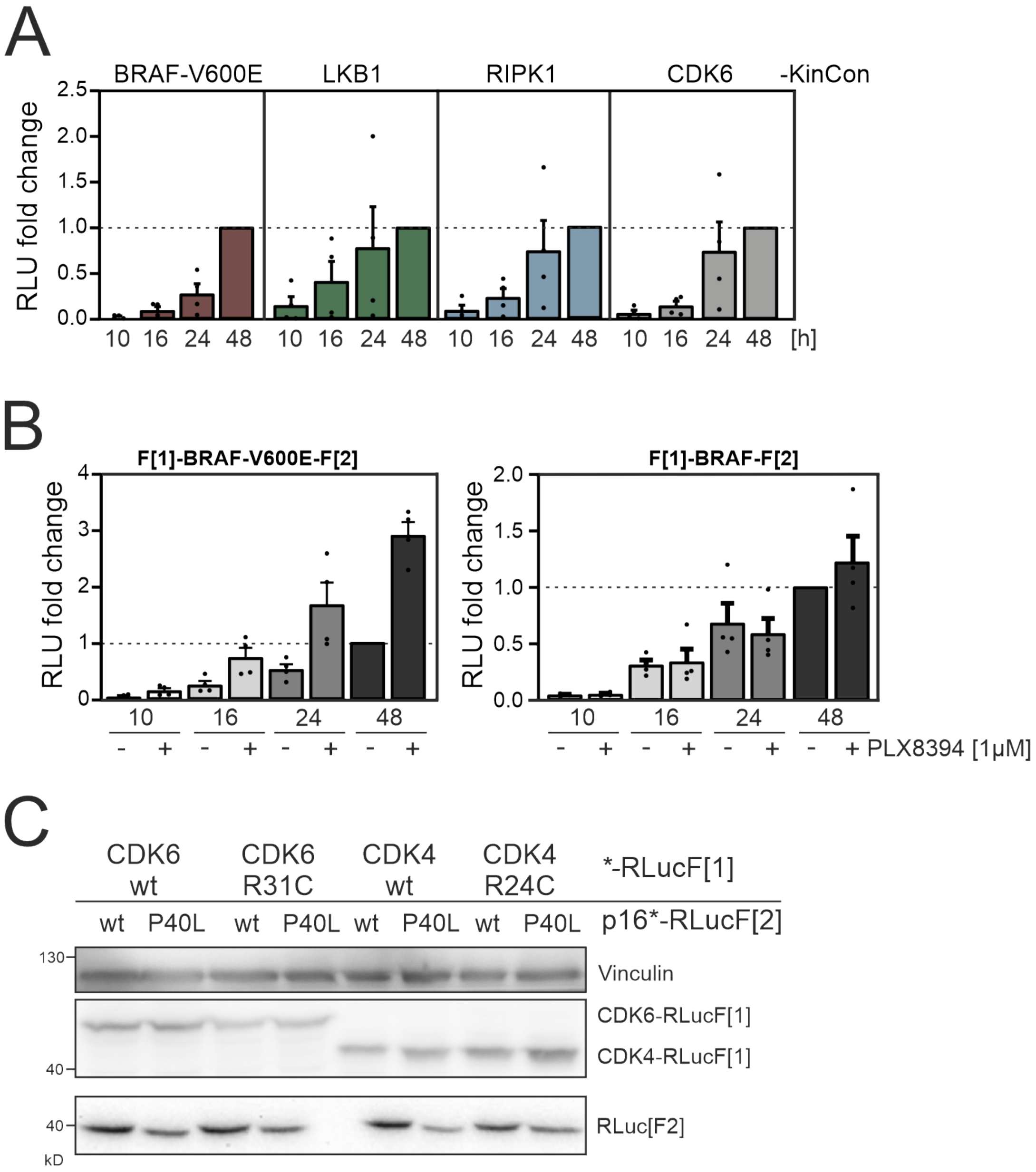


**Figure 1.** Time-dependent KinCon expression and CDK4/6:p16INK4a PPI **A)** Summary of n=4 independent experiments of time-dependent expressions of indicated KinCon reporter constructs in HEK293T cells is shown (mean ± SEM). BRAF-V600E, LKB1, RIPK1 and CDK6 KinCon reporters were transiently over-expressed in 24-well format in HEK293T cells for 10h, 16h, 24h and 48h each. **B)** Impact of 1μM PLX8394 exposure for 1h on BRAF and BRAF-V600E KinCon reporters transiently over-expressed for indicated time frames in HEK293T cells is shown. Mean of n=4 independent experiments with ±SEM is presented. **C)** Exemplary western blot of CDK4/6 – p16INK4a PPI PCA. Immunoblot with respective antibodies show expression levels of RLuc[F1] and RLuc[F2] tagged expression constructs.

#### Figure Supplement 2

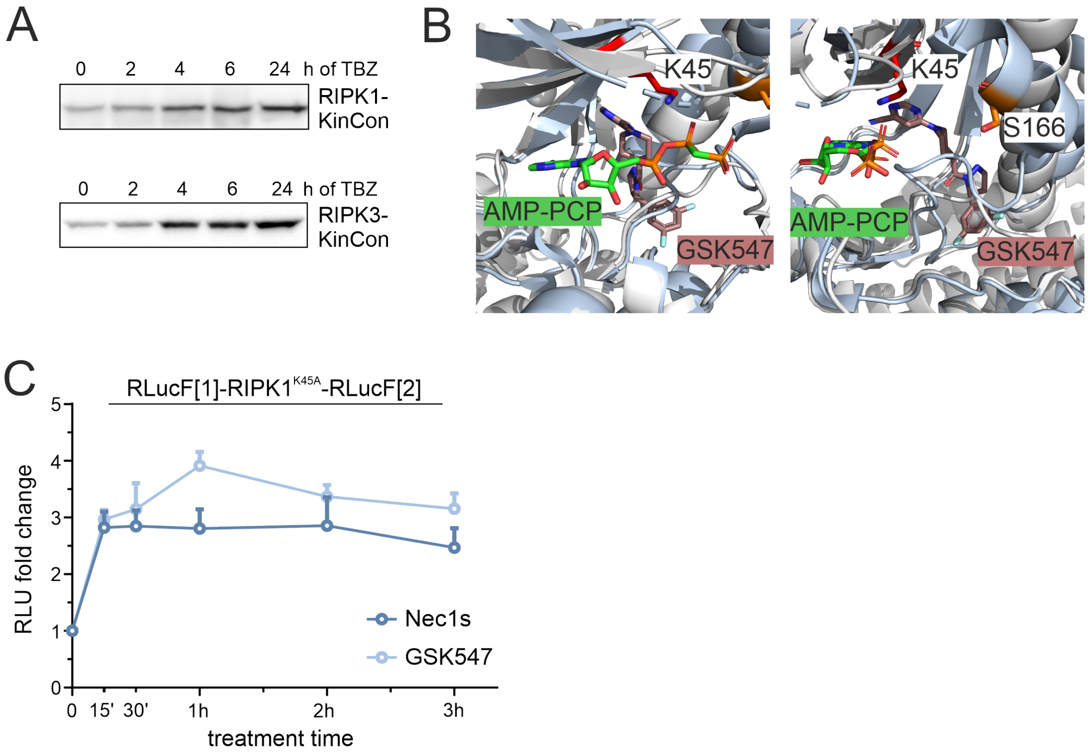


**Figure 2.** Small molecule effects on RIPK1. **A)** Representative western blots of RIPK1 and RIPK3 KinCon expressions after TBZ treatment for indicated time points. HEK293T cells were transfected with KinCon constructs and subjected to Bioluminescence measurements after 48h. **B)** 3D structure of RIPK1 with the inhibitor GSK547 (PDB code: 6HHO (***Wang et al.*** (***2018***))), which binds to an allosteric site in close proximity to ATP. For comparison, the structure of RIPK2 (light grey) in complex with the ATP analogue AMP-PCP (green sticks) is aligned (PDB code 5NG0 ***Pellegrini et al.*** (***2017***) C) Time dependent treatment of RIPK1-K45A KinCon with two RIPK1i (GSK547 and Necrostatin 1µM). The KinCon reporter was over-expressed in HEK293T RIPK1 KO cells for 48h. Points represent the mean RLU fold change relative to time point zero for n=3 independent experiments with ±SEM.

#### Figure Supplement 3

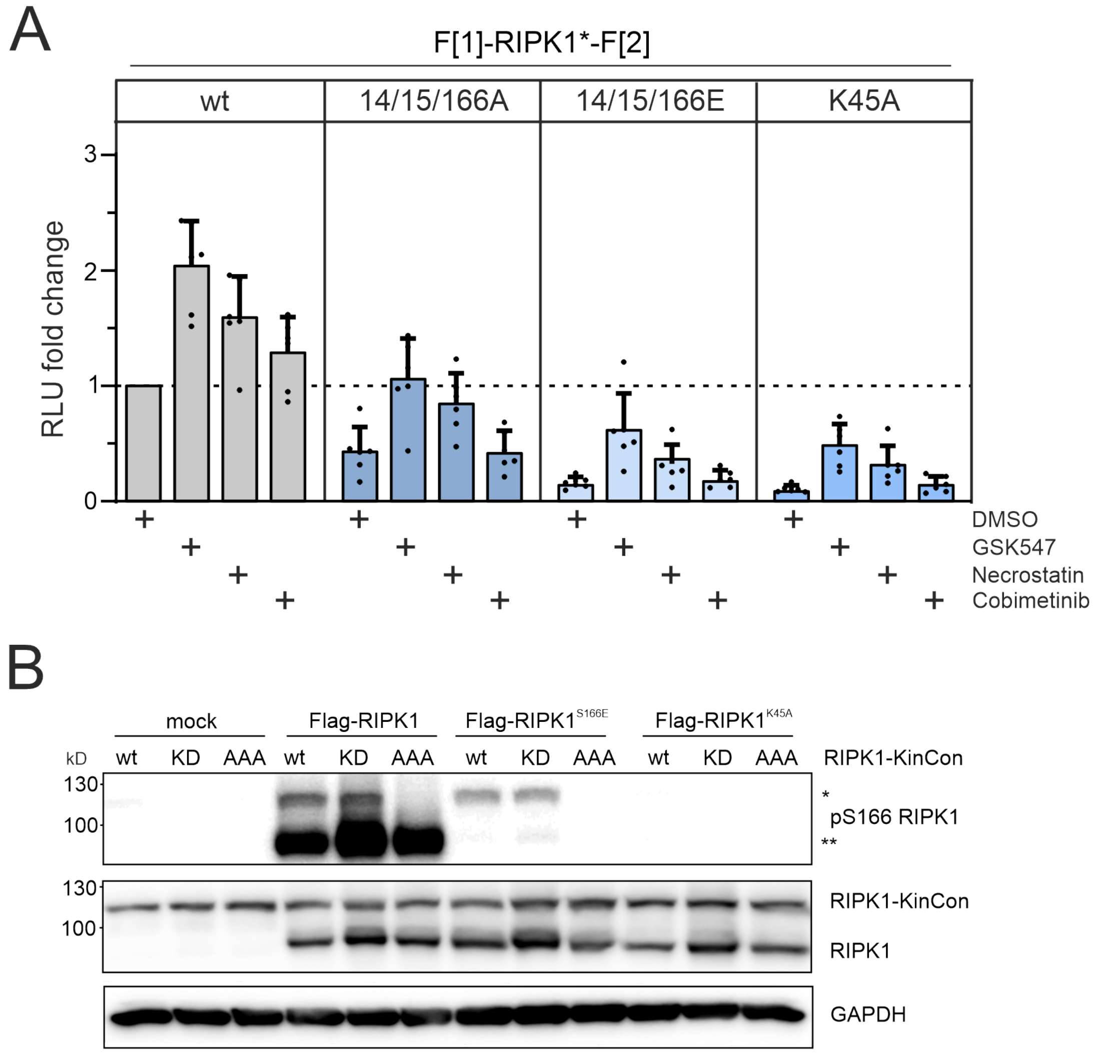


**Figure 3.** Conformation and phosphorylation of RIPK1. **A)** Dynamics of RIPK1 mutations upon exposure to the two RIPK1i (GSK547 and Necrostatin 1µM), and the MEKi Cobimetinib (1µM, control experiment) or DMSO for 1h. Bars represent the RLU fold change relative to the DMSO control of wt RIPK1 (mean ±SEM, n=6 ind. experiments, HEK293T RIPK1 KO). **B)** RIPK1 KinCon auto-phosphorylation with co-expressed Flag-tagged RIPK1 constructs following expression in HEK293T RIPK1 KO cells. Indicated RIPK1 wt or mutant constructs were co-expressed for 48h and subjected to western blotting. KinCon RIPK1 constructs are annotated with single asterisk and Flag-tagged RIPK1 with double asterisks.

### Figure Supplement 4

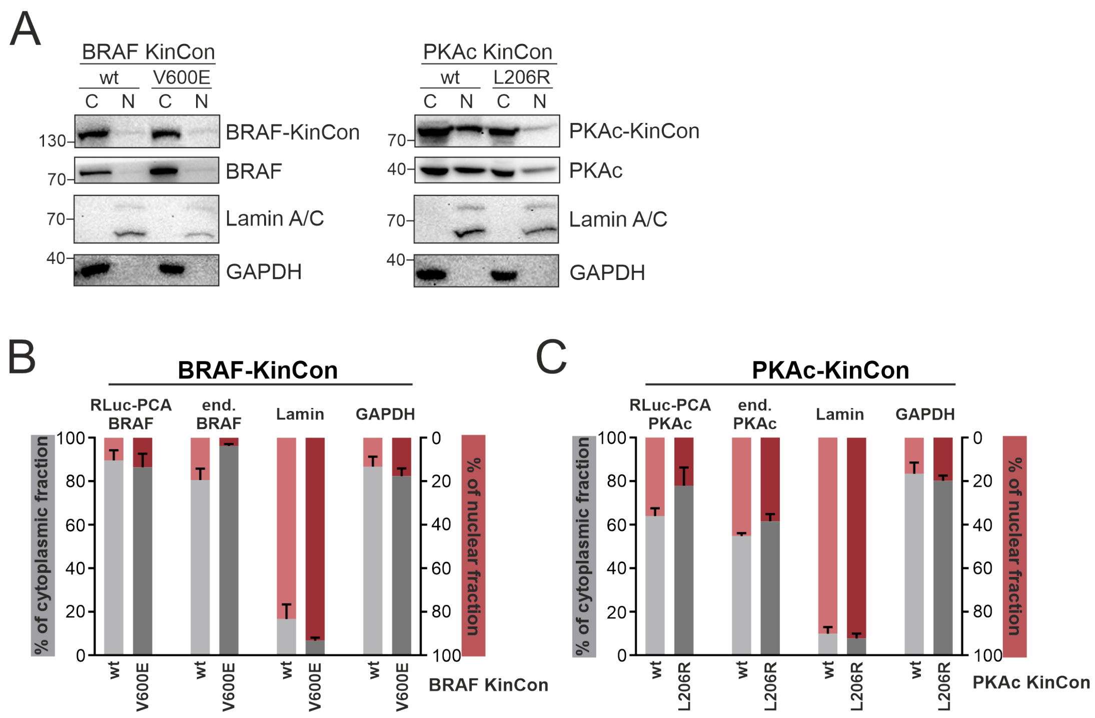


**Figure 4.** Subcellular Localization of KinCon reporters. BRAF and PKAc wt and mutated (BRAF V600E, PKAc L206R) reporters were overexpressed in HEK293T cells. **A)** Representative western blot from the cytoplasmic (C) and nuclear (N) fractions. **B+C)** Quantifications of the signals from n=3 ind. experiments (mean ±SEM).

**Figure Supplement 5**

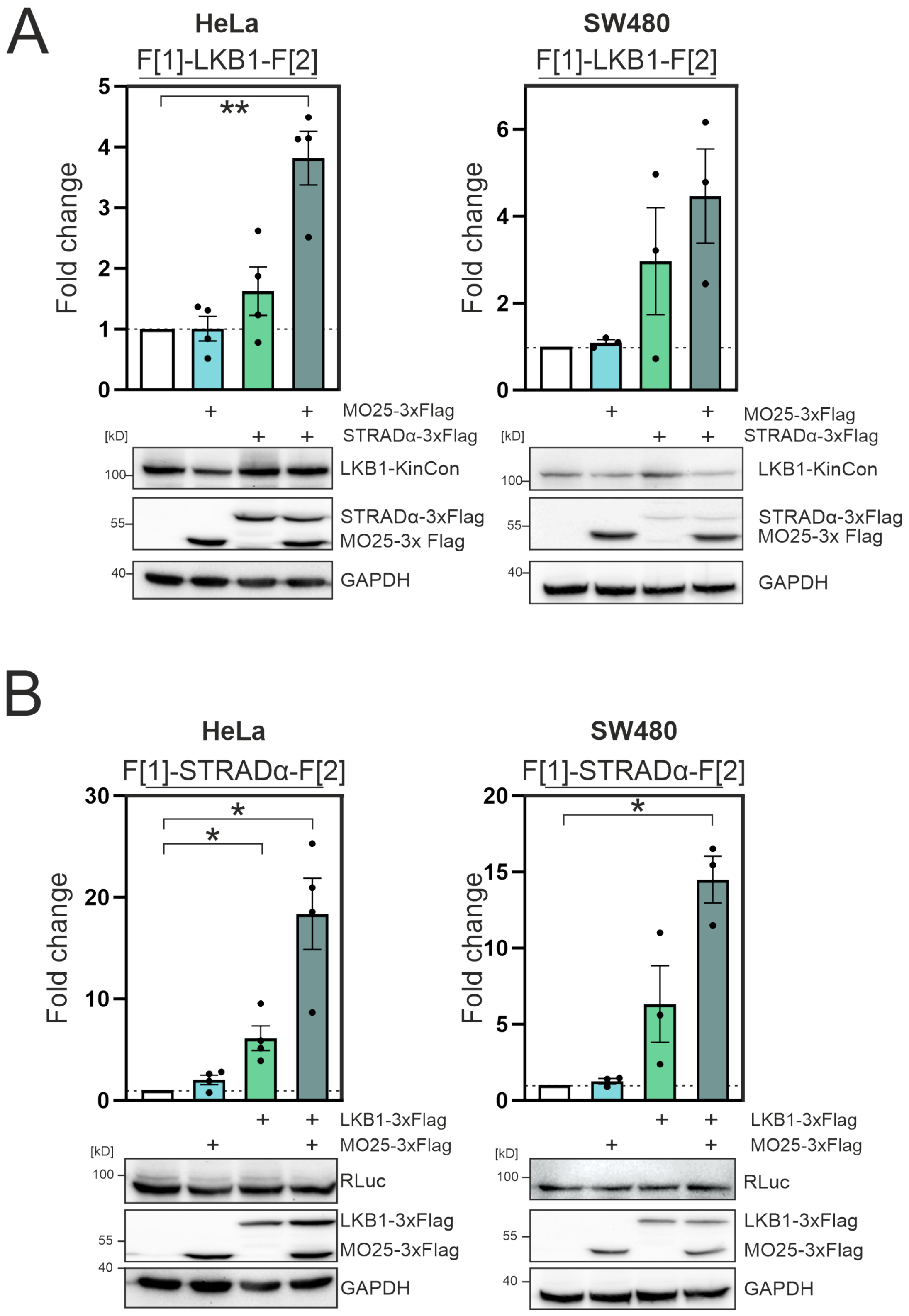

**Figure 5.** Complex formation of LKB1/STRADa/MO25 in HeLa and SW480 cells **A)** Effect of LKB1-STRAD$\alpha$-MO25 complex formation on the LKB1 KinCon reporter (HeLa and SW480 cells). Bioluminescence signals were compared to the LKB1-KinCon signal (mean $\pm$SEM, n=3-4). Representative western blots are shown below. **B)** Effect of LKB1-STRAD$\alpha$-MO25 complex formation on the STRAD$\alpha$ KinCon reporter (HeLa and SW480 cells). Bioluminescence signals were compared to the STRAD$\alpha$ -KinCon signal (mean $\pm$SEM, n=4 (HeLa) and n=3 (SW480)). Representative western blots are shown below. Statistical significance for A and B: One-sample t-test (*p<0.05, **p<0.01, ***p<0.001).
